# Coordinated morphogenesis through tension-induced planar polarity

**DOI:** 10.1101/207209

**Authors:** Ghislain Gillard, Ophélie Nicolle, Thibault Brugières, Sylvain Prigent, Mathieu Pinot, Grégoire Michaux

**Author notes:** Current address: MRC LMB, Cambridge, UK.

## Abstract

Tissues from different developmental origins must interact to achieve coordinated morphogenesis at the level of a whole organism. *C. elegans* embryonic elongation is controlled by actomyosin dynamics which trigger cell shape changes in the epidermis and by muscle contractions, but how the two processes are coordinated is not known. We found that a tissue-wide tension generated by muscle contractions and relayed by tendon-like hemidesmosomes in the dorso-ventral epidermis is required to establish a planar polarity of the apical PAR module in the lateral epidermis. This planar polarized PAR module then controls actin planar organization, thus determining the orientation of cell shape changes and the elongation axis of the whole embryo. This trans-tissular mechanotransduction pathway thus contributes to coordinate the morphogenesis of three embryonic tissues.

---

Understanding how tissues from different developmental origins interact to achieve coordinated morphogenesis at the level of a whole organism has been mostly studied through the prism of biochemical signaling pathways controlling the activity of transcription factors where one tissue sends a chemical signal received by another tissue (Hubaud and Pourquie, 2014; Petit et al., 2017; Schmid and Hajnal, 2015). However morphogenesis of epithelial tissues can also be controlled by biomechanical pathways physically linking two tissues (Aigouy et al., 2010; Butler et al., 2009; Collinet et al., 2015; Olguin et al., 2011; Sagner et al., 2012). The morphogenetic step of *C. elegans* embryonic elongation requires the direct coordination of three tissues: muscles, dorso-ventral epidermis and lateral epidermis (See Fig. S1A-D for a schematic representation of *C. elegans* anatomy). As was shown in a pioneering study, a mechanical signal generated by muscle contractions in the antero-posterior (A/P) axis from mid-elongation is translated into a biochemical pathway to control the maturation of tendon-like structures called *C. elegans* hemidesmosomes (CeHDs; Fig. S1D) that link muscles to the apical surface of the dorso-ventral epidermis (Zhang et al., 2011). However elongation starts by cell shape changes in the lateral epidermis (Costa et al., 1998; Vuong-Brender et al., 2016) where actomyosin contractions generating a tension in the dorso-ventral (D/V) axis (Vuong-Brender et al., 2017) are essential to initiate and maintain embryonic elongation. How the tissue-scale forces generated by muscles in the A/P axis and cell-scale forces generated by actomyosin contractions in the D/V axis are coordinated is not known and we decided to investigate the mechanisms underlying this coordination.

As was recently reported (Vuong-Brender et al., 2017) actin becomes progressively planar polarized in the D/V axis in lateral cells during elongation. It is partially disorganized at the 1.5-fold stage (Fig. 1A) whereas it becomes robustly oriented along the D/V axis from the 2fold stage (Fig. 1B-C). We quantified this actin orientation as previously described (Fig. 1D-F) (Cetera et al., 2014). To identify new factors involved in actin planar polarity in the lateral epidermis we performed a small-scale RNAi screen targeting candidate genes implicated in epithelial junctions, polarity, cytoskeleton and membrane traffic (Supplementary Table 1) which identified six genes. The first four genes are involved in muscle function or CeHDs formation (Fig. S1D) and their depletion lead to the Pat (for Paralyzed and Arrested at the Two-fold stage) phenotype. Two of these genes are expressed in muscles: *unc-112* (Fig. 1H, J), a kindlin homologue essential for myofilament assembly and localization of the PAT-3 β-integrin (Rogalski et al., 2000), and *pat-4* (Fig. S1F, H), a muscle kinase homologous to mammalian ILK (Mackinnon et al., 2002). The other two genes are expressed in the D/V epidermis: *unc-52* (Fig. S1G, I), encoding an extracellular matrix (ECM) perlecan secreted basolateraly (Rogalski et al., 1993) and *vab-10* (Fig. 1I, K), a spectraplakin homologue and a structural protein of CeHDs (Bosher et al., 2003). Given the known function of these genes and the function of CeHDs in relaying muscle contractions to the D/V epidermis (Zhang et al., 2011) it surprising that they control actin planar polarity in the lateral epidermis where none of them is expressed; we concluded that the mechanical signal initiated by muscle contractions is likely transmitted to the lateral epidermis via the D/V epidermis.

**Figure 1:**
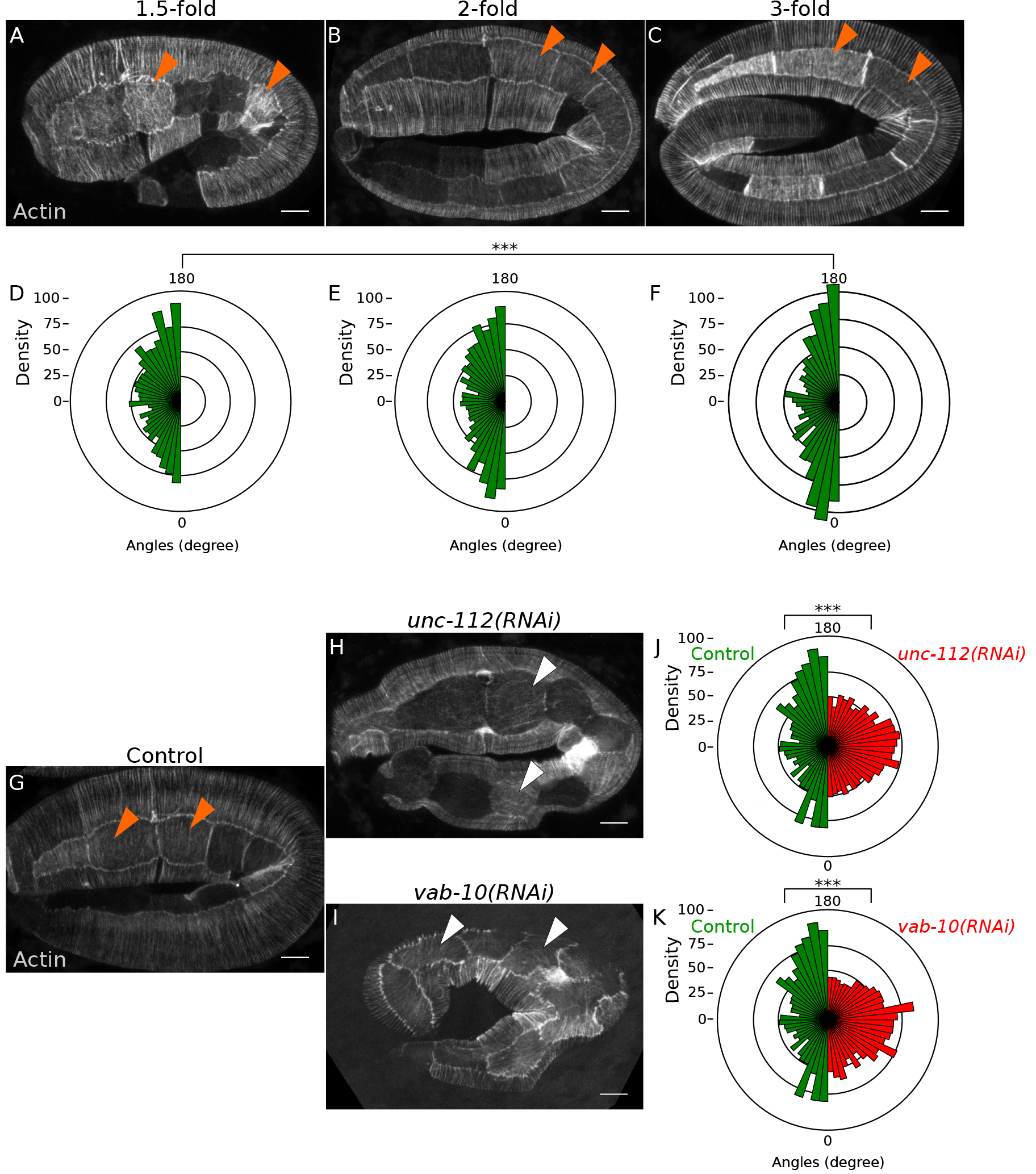
Muscles and CeHDs are required for actin planar polarity. **A-C** Actin orients along the dorso-ventral axis during elongation, from the 1.5-fold stage, where it exhibits a partial orientation along the D/V axis (n=19 embryos, 51 cells) to the 2-fold (n=18 embryos, 57 cells) and the 3-fold stages (n=15 embryos, 48 cells) where actin is strongly oriented along the D/V axis (orange arrowheads). **D-F** Actin orientation in epidermal cells has been quantified for each cell as described (Cetera et al., 2014). Graphs represent the distribution of angles compared to the D/V axis. **G-I** Actin organization in 2-fold stages embryos; actin is disorganized in epidermal cell under *unc-112* (n=17 embryos, 61 cells) and vab-10 (n=17 embryos, 39 cells) depletion. Orange arrowheads indicate proper actin polarization in control cells (n=11 embryos, 34 cells) while white arrowheads indicate cells where actin is strongly disorganized. **J-K** Quantification of actin orientation under *unc-112* and *vab-10* depletion, respectively. Scale bars: 5 μm.

Our screen next identified an unexpected role for the small GTPase RAB-1, a protein well described for its role in membrane traffic between the ER and the Golgi in yeast and mammalian cells (Plutner et al., 1991; Segev et al., 1988). We found that *rab-1* depletion leads to a 2-fold arrest (Fig. S2A-B) and a complete actin disorganization reminiscent to that observed under CeHDs depletion (Fig. 2A-C). *rab-1(RNAi)* specificity was established in a strain expressing an RNAi-insensitive *rab-1* version from *C. briggsae* and the presumptive null allele *rab-1(ok3750)* induces an elongation arrest similar to *rab-1(RNAi)* treatment (Fig. S2A-B). RAB-1 is ubiquitously expressed and its localization in epidermal cells is consistent with an ER-Golgi function (Fig. S2C). In *C. elegans* RAB-1 has been implicated in the secretion of the yolk receptor RME-2 (Balklava et al., 2007) and we reasoned that RAB-1 could have a function in regulating CeHDs remodeling as was shown for calreticulin, an ER enzyme (Zahreddine et al., 2010); we therefore examined the endogenous localization of the secreted UNC-52 perlecan and the basolateral transmembrane receptor LET-805 (Hresko et al., 1999), both synthesized in the dorso-ventral epidermis. We found that *rab-1* depletion leads to a disruption of UNC-52 and LET-805 localization (Fig. 2D-G). These observations, together with results presented below, strongly suggest that the actin defect observed in the lateral epidermis under *rab-1* depletion is the consequence of CeHDs disruption.

**Figure 2:**
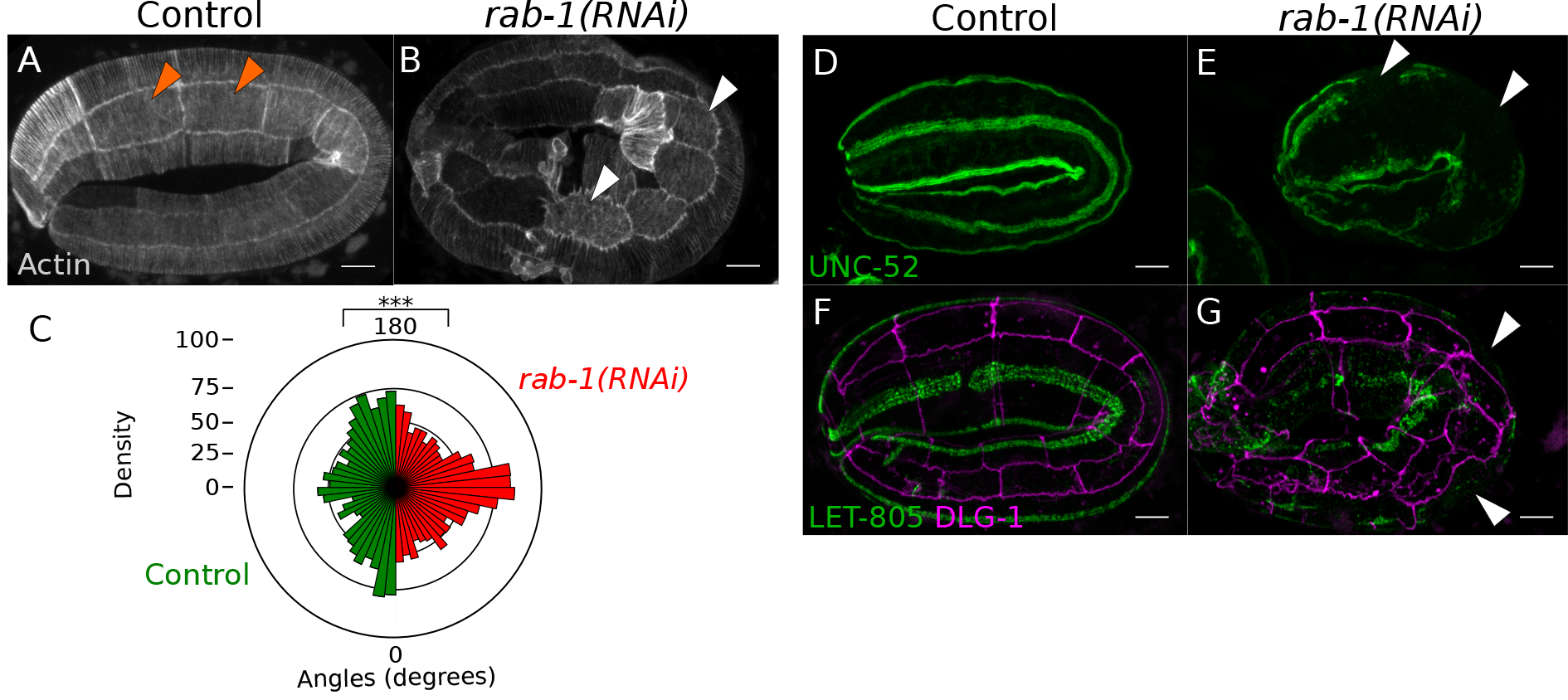
RAB-1 controls actin orientation in the lateral epidermis through CeHDs components trafficking. **A-C** *rab-1* depletion strongly disorganizes actin in lateral epidermal cells. Orange arrowheads indicate proper actin polarization in control cells (n=16 embryos, 54 cells) while white arrowheads indicate cells where actin is disorganized under *rab-1* depletion (n=16 embryos, 69 cells). **D-G** *rab-1* depletion disrupts the localization of UNC-52 as revealed by immunostaining (D, E; n=20 control embryos, n=15 *rab-1(RNAi)* embryos) and of LET-805 as assessed by endogenous localization in a CRISPR strain (F, G; n=30 control embryos, n=24 *rab-1(RNAi)* embryos). White arrowheads indicate areas of interrupted staining; DLG-1 is a junction marker. All embryos are imaged at the 2-fold stage. Scale bars: 5 μm.

The last candidate gene identified in our screen encodes the essential polarity protein PAR-3 which also controls actin organization in lateral cells (Fig. 3A-C). Surprisingly the localization of the PAR-3/PAR-6/PKC-3 polarity module has never been described in the epidermis beyond early elongation, either by immunostaining or using reporter constructs. Using CRISPR/Cas9 genome edited strains to localize endogenous PAR-3, PAR-6 and PKC-3 we found that they all exhibit planar polarity in lateral cells, accumulating preferentially on junctions between lateral cells (L-L junctions) but not on junctions between cells of the lateral and dorso-ventral epidermis (L-D/V junctions) (Fig. 3D-G, L-L', O-O' and Fig S3A-F”); we did not observe PAR-3 recruitment to the junctions between ventral or dorsal cells which are parallel to the L-L junctions. We found that PAR-3 accumulates progressively at the L-L junctions (Fig. 3D-G, Fig. S3A-C”; a similar observation was made for PAR-6 (Fig. S3D-F”) and PKC-3). Based on its planar polarized localization restricted to lateral cells, the PAR module seems to be required specifically in these cells. We therefore decided to test whether the PAR planar polarity was dependent on muscle contractions and dorsoventral CeHDs as we showed for actin. We found that the depletion of genes controlling muscle contractions or CeHD components, identified previously as required for correct actin planar polarity, leads to a loss of PAR-3 accumulation at the plasma membrane (Fig. 3H-K and Fig. S3G-J’). The planar polarity of PAR-6 and PKC-3 is also affected under RAB-1 depletion (Fig. S3M-R). To test the requirement of the whole apical PAR module we showed that PAR-3 is essential to properly recruit PAR-6 and PKC-3 at L-L junctions (Fig. 3L-Q), while PAR-6 enables the recruitment of PAR-3 (Fig. S3K-L’). We concluded that there is a co-dependency between the members of the PAR module for their planar polarized localization and that the PAR module is needed to properly orient actin in the D/V axis in the lateral epidermis.

**Figure 3:**
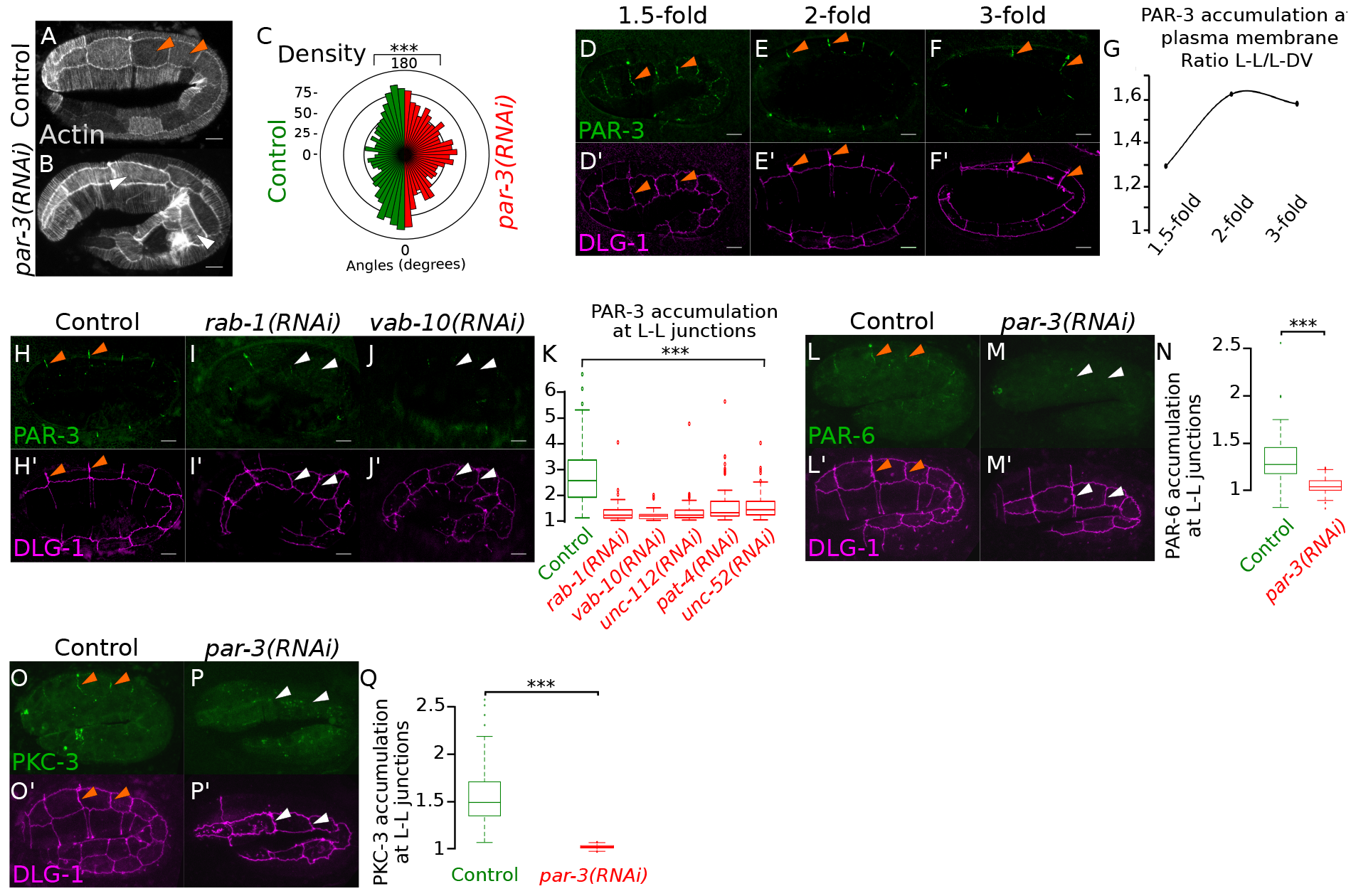
Muscles and CeHDs control PAR planar polarity in the lateral epidermis. **A-C** *par-3* depletion leads to actin misorientation (white arrowheads in B, n=15 embryos, 67 cells, compared to orange arrowheads in control, n=11 embryos, 34 cells) as revealed by the associated quantification in C. **D-G** Endogenous PAR-3::GFP accumulates at L-L junctions during elongation. Orange arrowheads indicate L-L junctions where PAR-3 tends to accumulate during elongation. DLG-1 (purple) corresponds to junctions. Quantification was made by measuring the ratio between PAR-3 staining at L-L and L-D/V junction for each stage (G, n=10 embryos 1,5-fold, 79 L-L / 137 L-D/V; n=10 embryos 2-fold, 80 L-L / 113 L-D/V; n=10 embryos 3-fold, 68 L-L / 113 L-D/V). **H-K** Depletion of *rab-1* or *vab-10* by RNAi leads to an absence of recruitment of endogenous PAR-3::GFP at L-L junctions as depicted by white arrowheads compared to orange arrowheads in control. This absence of PAR-3 localization at the plasma membrane has been quantified in **K** (n=29 control embryos, 169 LL; n=12 embryos *rab-1(RNAi)*, 61 L-L; n=13 embryos *vab-10(RNAi)*, 82 L-L, n=29 embryos *unc-112(RNAi)*, 188 L-L; n=19 embryos *pat-4(RNAi)*, 132 L-L; n=21 embryos *unc-52(RNAi)*, 106 L-L). **L-Q** PAR-3 is required for the recruitment of endogenous PAR-6::GFP (L-N) and GFP::PKC-3 (O-Q) at L-L junctions. N and Q correspond to quantifications of PAR-6 and PKC-3 recruitment, respectively. All embryos are imaged at the 2-fold stage except in panels D and F. Scale bars: 5 μm.

How could the PAR module and the actin cytoskeleton be organized in the lateral epidermis from the 2-fold stage by muscle contractions without direct connections between muscles and lateral epidermal cells? One obvious candidate corresponds to the E-cadherin-containing adherens junctions (AJs) between the dorso-ventral and lateral epidermal cells. We therefore first examined the possibility that muscle contractions and CeHDs could control AJs organization or the global epithelial polarity of lateral cells. To test this hypothesis we examined the impact of *rab-1* depletion which could by itself control E-cadherin (E-cad) localization and/or dynamics. In *C. elegans* epidermal cells AJs are located at the apical side above the diffusion barrier composed by the DLG-1/AJM-1 complex (Gillard et al., 2015). As revealed by the localization of a functional E-cad::GFP (Achilleos et al., 2010) these junctions seem unaffected following *rab-1* depletion by RNAi (Fig. 4A-B) or in a *rab-1* deletion allele (Fig. S4A-B). Moreover the weak planar polarity of E-cad is not altered as shown by the quantification of the ratio of E-cad at L-D/V junctions versus L-L junctions (Fig. 4C) despite a slight decrease in overall E-cad membrane accumulation (Fig. 4D) consistent with the canonical RAB-1 function in secretion. Similarly, the localization of VAB-9, another component of apical AJs in *C. elegans (Simske et al., 2003)*, is not affected (Fig. S4C-D). However AJs remodeling is essential to promote morphogenesis in the Drosophila embryo (Levayer and Lecuit, 2013; Rauzi et al., 2010) and we also examined the E-cad dynamics by fluorescence recovery after photo-bleaching (FRAP) experiments. We did not identify a significant effect of *rab-1* depletion on E-cad dynamics (Fig. 4E-G) and the immobile fraction of E-cad which reaches around 60% is consistent with previous reports (Bulgakova et al., 2013; Erami et al., 2015). This junctional integrity was confirmed by electron microscopy (Fig. S4E-F). To test the integrity of the diffusion barrier and of apico-basal polarity we established that the localization of the apical transmembrane protein CHE-14 (Gillard et al., 2015; Michaux et al., 2000) and the basolateral polarity determinant LET-413/Scribble (Legouis et al., 2000) were not affected in the absence of RAB-1 (Fig. S4G-J). Finally we did not identify a genetic interaction between the α-catenin hypomorphic allele *hmp-1(fe4)* (Simske et al., 2003) and *rab-1(RNAi)* (Fig. S4K). Altogether these results demonstrate that junction integrity and the apico-basal polarity of lateral cells is not dependent on muscle contractions and CeHDs. However these results do not imply that AJs are not implicated in the transduction of the mechanical signal generated by muscles between the dorso-ventral epidermis and the lateral epidermis. To directly test this possibility and because the loss of function of HMP-1/α-catenin leads to an arrest around the 1.5-fold stage, we depleted *hmp-1* by RNAi. We found that PAR-3 fails to be recruited to the L-L junctions in *hmp-1(RNAi)* embryos (Fig. 4H-J). We therefore propose that the mechanical signal initiated by muscles and CeHDs in the dorso-ventral epidermis is relayed to the lateral epidermis via AJs.

**Figure 4:**
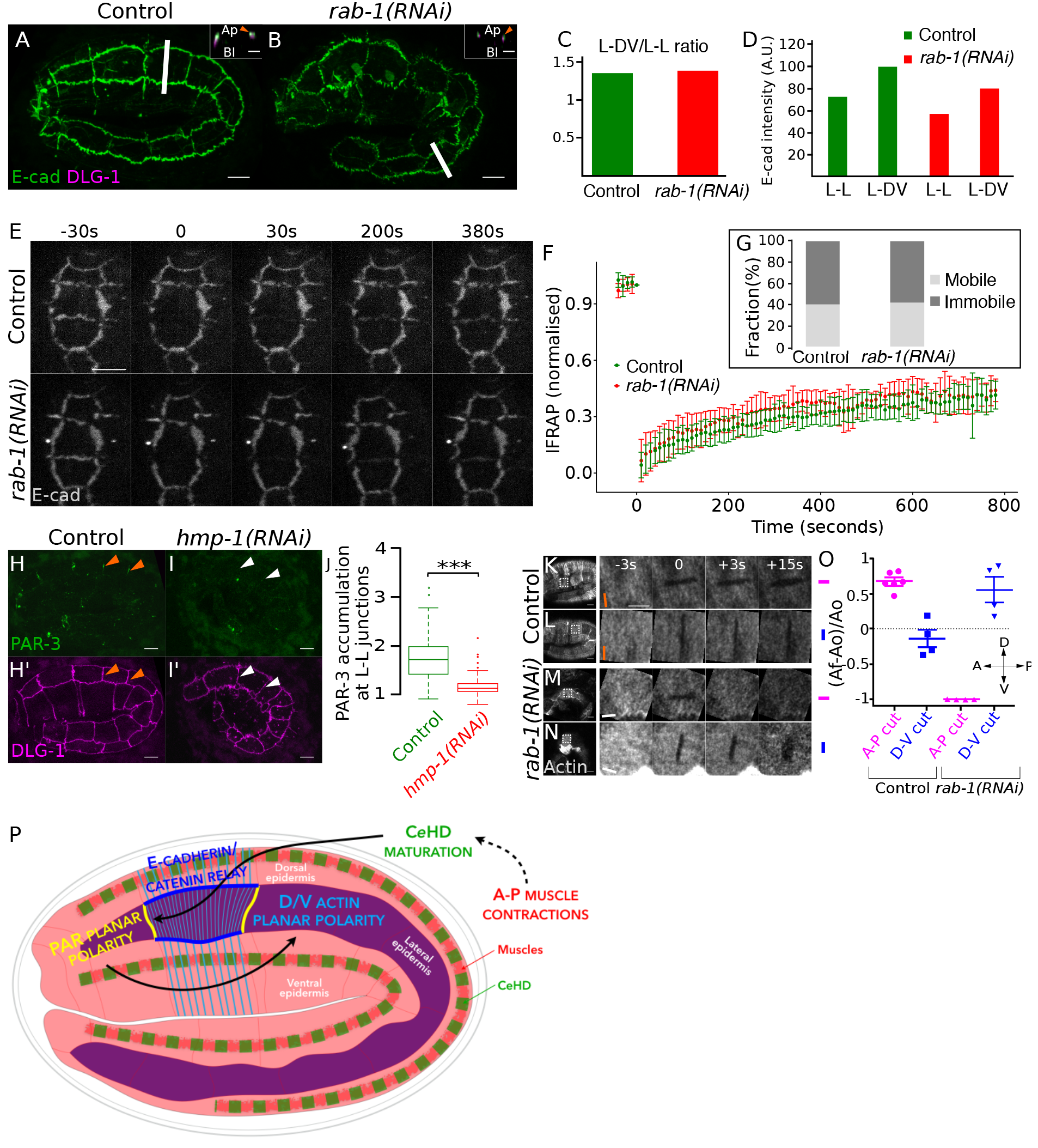
AJ function in signal transduction. **A-D** E-cad remains apical, above DLG-1 upon *rab-1* depletion (B, n=29 embryos) as in control embryos (A, n=18). Small insets correspond to Z-section represented by a white line in the associated picture. The quantification shows that the ratio of E-cad between L-D/V and L-L junctions remains intact under *rab-1* depletion (C). However, there is a slight decrease in E-cad overall accumulation at plasma membrane under *rab-1* depletion (D). **E-G** FRAP experiments performed in 1.5-fold stage embryos show that E-cad dynamics is not affected in the absence of RAB-1 (n=7 junctions for control, n=8 for RNAi embryos), as depicted on the recovery curves (F) and the percentages of mobile and immobile fractions (G). **H-J** Endogenous PAR-3::GFP localization at L-L junctions observed in control embryos (orange arrowheads, n=26 embryos, 78 junctions) is lost under *hmp-1* depletion (white arrowheads, n=37 embryos, 111 junctions). J corresponds to quantifications of PAR-3 recruitment. **K-O** Nano-ablation experiments of the actin cytoskeleton in the lateral epidermis in control (K, L, actin oriented in the D-V axis as shown by the orange bar) and *rab-1*-depleted (M, N, actin oriented in the A-P axis as shown by the white bar) embryos. Cuts were performed along the A-P axis (K, M, in purple) or along the dorso-ventral axis (L, N, blue). These experiments have been quantified by measuring the relative expansion of the cut area over time (O). *rab-1* depletion affects tension forces orientation. All embryos are imaged at the 2-fold stage except in E. Scale bars: 5 μm, except for small insets: 2 μm. **P** Working model. Muscle contractions promote the assembly and the stabilization of CeHDs (Zhang et al., 2011) to enable force transmission between muscles and epidermis during morphogenesis. This force transmission in the dorso-ventral epidermis is required to promote planar polarization of lateral epidermal cells, most likely via the E-cad/catenin complex. This leads to PAR protein accumulation at L-L junctions and ultimately actin orientation along the dorso-ventral axis, a prerequisite to properly control cell shape changes and contribute to morphogenesis along the antero-posterior axis.

We finally wanted to assess the link between morphogenesis and actin disorganization in the lateral epidermis following CeHDs disruption. We therefore exploited a phenotype observed in some embryos where actin orientation was shifted by 90° and aligned along the A/P axis: this particular phenotype allowed us to test the hypothesis that the orientation of actin and of tension would be directly linked. We used laser nano-ablation experiments on the actin cytoskeleton to compare the orientation of tension forces in lateral epidermal cells depending on the orientation of actin itself. In a control situation, actin is oriented in the D/V axis, perpendicular to the elongation (A/P) axis. A cut in the A/P axis leads to a strong relaxation of the actin cytoskeleton, whereas a cut along the D/V axis does not lead to any relaxation (Fig. 4K, L, O). These results reveal that tension forces are oriented along the dorso-ventral axis in the lateral epidermis as was found in a recent study (Vuong-Brender et al., 2017). Conversely, in the cells of *rab-1* depleted embryos where actin is orientated in the A/P axis we found a corresponding 90° shift or a disorganization of tension forces (Fig. 4M-O). Because acto-myosin contractions in the lateral epidermis are required for embryonic elongation (Vuong-Brender et al., 2016), we postulate that this disorganization of tension forces is at least partly responsible for the arrest at the 2-fold stage observed under *rab-1* and CeHDs depletion.

We have identified a new trans-tissular signaling pathway required to coordinate morphogenesis between three tissues: muscles, the dorso-ventral epidermis and the lateral epidermis during *C. elegans* embryonic morphogenesis (see working model in Fig. 4P). We found that muscle contractions and force transmission through CeHDs and AJs are essential to promote a planar organization of the PAR module and of actin. Tissue mechanics and force transmission have previously been involved in drosophila wing morphogenesis where the wing hinge retraction enables PCP rearrangement from a radial to a proximal-distal-oriented polarity in the wing blade (Aigouy et al., 2010); similarly a mechanical input has been implicated in setting up planar polarity during drosophila germ-band extension (Butler et al., 2009; Collinet et al., 2015; Simoes Sde et al., 2010). In a different context a recent study demonstrated that a physical pressure exerted by proliferating dermal cells controls the patterning of the avian skin (Shyer et al., 2017). However in all these examples the tissue-wide tension is generated in parallel with the target tissue, while during *C. elegans* morphogenesis the tension generated by muscles is perpendicular to the local tension exerted by actin. Interestingly smooth muscles lying below or inside many tissues such as the skin, the intestine, the respiratory organs or the reproductive tracts could be essential for planar polarity establishment in surrounding epithelial tissues throughout the animal kingdom.

## ACKNOWLEDGEMENTS

We thank Aurélien Bidaud-Meynard, Michel Labouesse, Roland Le Borgne, Anne Pacquelet and Sophie Quintin for helpful discussions and critical reading of the manuscript and Yann Le Cunff for his help in statistical analysis. We are grateful to Liam Coyne, Anushae Syed, Ken Kemphues, Bob Goldstein and Michel Labouesse for sharing unpublished CRISPR/Cas9 genome edited GFP fusion strains, as well as Jeff Hardin, Renaud Legouis and Jeremy Nance for strains. We thank Maïté Carré-Pierrat and the UMS3421 (Lyon) for generating the *gfp::rab-1* expressing strain. Some strains were provided by the CGC, which is funded by NIH Office of Research Infrastructure Programs (P40 OD010440; University of Minnesota, USA). We thank the photon and electron microscopy facilities of the Microscopy Rennes Imaging Center. This work was supported by the Ligue contre le Cancer Grand Ouest [22/29/35/72], the Fondation ARC and the Fondation Maladies Rares; we also receive institutional funding from the Centre national de la recherche scientifique (CNRS) and the Université de Rennes 1. GG was founded by the Ligue Nationale Contre le Cancer (2015-2016).

## Author contributions

G.G. and G.M. designed the experiments. G.G., O.N., T.B. and G.M. performed experiments and data analysis. S.P. and M.P. helped to design some experiments and perform quantifications. G.G. and G.M. wrote the manuscript.

## MATERIAL AND METHODS

### Genetics

*C. elegans* strains were maintained and crossed as described (Brenner, 1974). The strains used in this study are shown in Supplementary Table 2.

### Plasmids construction and strains

The *rab-1::gfp* construct under the control of its own promoter was generated by inserting the *gfp* gene after the *rab-1* promoter in frame with the *rab-1* ATG followed by the *rab-1* UTR. The *C. briggsae* version of *rab-1* (called *Cbr-rab-1*) was amplified by PCR then cloned in the Gateway pDONR p221. Both constructs were injected at 5ng/μL together with the *rol-6(su1006)* marker at 100ng/μL in the *C. elegans* N2 strain.

### RNAi

Embryonic RNAi was performed by feeding as described using the Ahringer-Source BioScience library (Fire et al., 1998; Kamath and Ahringer, 2003; Shafaq-Zadah et al., 2012); RNAi was induced in young adults and the phenotypes observed in the next generation (F1). L4440 corresponds to the standard control RNAi feeding strain. For some genes (e.g. *par-3*, *par-6*) the duration of the RNAi treatment was adapted (<24h) to observe elongation phenotypes while avoiding earlier developmental phenotypes usually associated with these genes; *rab-1(RNAi)* was induced for about 20h to avoid the sterility triggered in the parental generation. All embryos observed and used for quantifications were 7-10h old, corresponding to 1.5-to 3-fold embryos in a WT strain; as a control we checked that there was not expression of the *myo-2p::GFP* transgene which is expressed from the 3-fold stage and present in the FL311 strain used to localise ABD::GFP. RNAi efficiency was checked by observing the induction of a developmental arrest whenever such a phenotype was expected based on previous reports. To test the specificity of the *rab-1(RNAi)* we scored the embryonic lethality observed following *Cel-rab-1(RNAi)* in an N2 strain and in a strain expressing *Cbr-rab-1* (Fig. S2a-b).

### Immunostaining

Fixation of embryos was performed as described using the freeze-crack methanol protocol (Leung et al., 1999). We used the anti-UNC-52 MH2 (1/50) monoclonal antibodies from DSHB (University of Iowa, USA). Alexa Fluor 488 antibody (Invitrogen) was used as secondary antibodies.

### Electron microscopy

Electron microscopy experiments were performed as described (Shafaq-Zadah et al., 2012). Briefly embryos were fixed by high pressure freezing, followed by freeze substitution, flat embedding to allow antero-posterior orientation and sectioning. control (n=3) and *rab-1(RNAi)* (n=4) embryos were observed. Each embryo was sectioned every 5-7μm to ensure that different cells were observed in different 5-7μm segments; 3 segments were examined for each embryo. Observations were performed on a Jeol JEM1400 equipped with a Gatan Orius SC1000 camera.

### Confocal microscopy and signal quantifications

Confocal observations were performed using Leica (Wetzlar, Germany) SPE, SP5 or SP8 confocals equipped with 63X objectives (LAS AF software). The SP5 and SP8 confocals are equipped with hybrid detectors which were used to image the low signals generated by the genome-edited strains expressing PAR-3::GFP, PAR-6::GFP and GFP::PKC-3 at the endogenous level. They were also used to image ABD::GFP at the highest possible resolution with a low background. All images were examined using ImageJ 1.43 and assembled using the Inkscape software.

The percentage of embryos displaying a phenotype was obtained either by direct observation or after quantification. Quantifications were performed using ImageJ 1.43 along straight lines (length 10μm, width 0.3μm) over the membrane and cytoplasmic parts of at least three cells for each embryo. The membrane quantification was normalized to the cytoplasmic background; a ratio of 1 therefore indicates no specific membrane staining.

Actin orientation was measured in Matlab using custom-routines developed elsewhere (Cetera et al., 2014). Briefly local directors representing actin alignment were determined as follows for each cell: a cell was properly oriented and broken in small overlapping windows of 2.6*2.6 mm and the 2D FFT of each filtered window was calculated, giving a range of angles whose values are given compared to the dorso-ventral axis.

### FRAP experiments and quantifications

FRAP experiments were performed on an inverted Nikon Ti-E microscope equipped with a Spinning-disk CSU-X1 and a FRAP head. Embryos were imaged with a 63X/1.4 PLAN APO objective and fluorescence was collected with a sCMOS ORCA Flash 4.0 camera. The FRAP was performed on a whole junction with 100% laser power, 50 iterations and a line thickness of 2, in the iLAS software in Metamorph. Post-FRAP images were acquired every 10 seconds. Quantifications were made manually in Image J by measuring the mean intensity of the bleached junction after background subtraction.

### Nano-laser ablations and quantifications

Nano-laser ablation experiments were performed on an inverted Leica SP5 microscope equipped with a Pulsed laser Mai Tai HP Ti. Embryos were imaged with a 63X/1.4 HCX PL APO objective. Nano-laser ablations were performed at 800 nm with a single iteration. Images were acquired every 1.27 seconds. For quantifications, the cut zone was manually tracked over time in ImageJ and the relative expansion was calculated as follow: (A_f_ − A_0_)/A_0_. A value of 1 thus indicates a relaxation, while 0 indicates no relaxation and -1 corresponds to a recovery of fluorescence in the cut zone.

### Statistical analysis

Parametric T-tests were used when samples had a Gaussian distribution and similar variances. Other cases were treated using non-parametric Wilcoxon tests. Significance is indicated as follow: * p<0.05, ** p<0.01, *** p<0.001.

---

**Figure S1:**
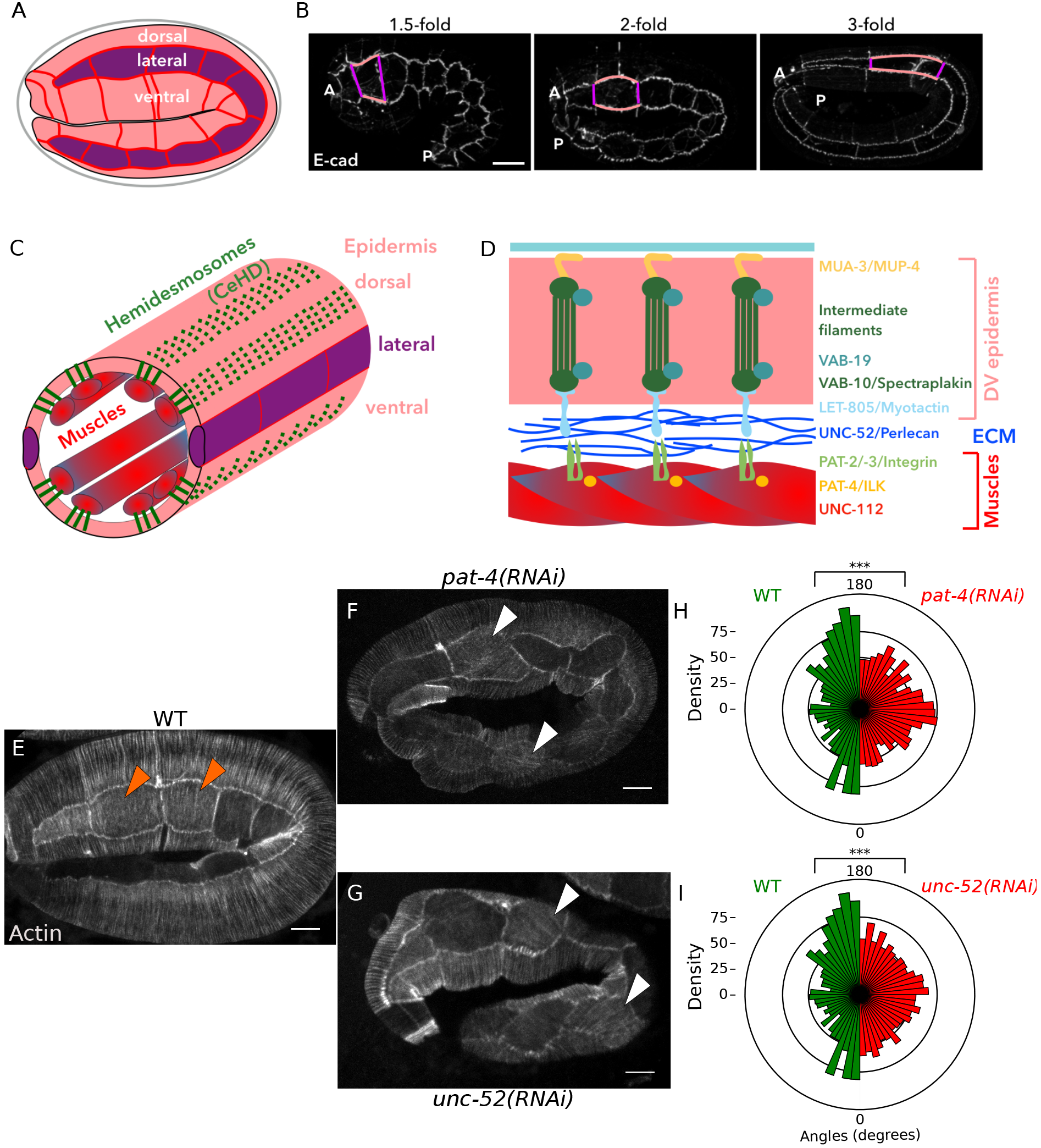
Muscles and CeHDs are required for actin planar polarity. **A-D** Schematic representation of *C. elegans* anatomy and junctional organization during morphogenesis. The epidermis is divided into three parts: two syncytia, the ventral and the dorsal epidermis in pink, and two rows of ten cells, one on each side of the embryo, the lateral epidermis, in purple (A); the dorsal and ventral epidermis have a similar developmental program and are referred to as the dorso-ventral epidermis. The elongation of the embryo reflects the elongation of the cells from the lateral epidermis. These cells exhibit a planar polarity in the sense that the “vertical junctions” (referred as L-L junctions, for lateral-lateral junctions, in purple) shrink during elongation while the “horizontal junctions” (referred as L-DV junctions, for lateral-dorsoventral junctions, in orange) elongate in the axis of embryonic elongation (B). The embryo can be assimilated to a tube surrounded by the epidermis. Muscles lay beneath the basolateral membrane of the dorso-ventral epidermis, linked to the apical side by hemidesmosome-like structures called CeHDs (for *C. elegans* hemidesmosomes, in green) (C). These CeHDs link the muscles to the dorso-ventral epidermis through the extracellular matrix (ECM) and the whole dorso-ventral epidermis thickness to the apical surface and the apical ECM. CeHDs are mainly composed by apical transmembrane receptors (MUA-3, MUP-4), intermediate filaments and linker proteins (VAB-19, VAB-10), a basolateral receptor (LET-805) and ECM protein UNC-52. The ECM is in turn linked to muscles through the action of integrins which require PAT-4 and UNC-112 for their proper localization (D). **E-G** Actin is disorganized in lateral epidermal cell under *pat-4* (n=19 embryos, 72 cells) and *unc-52* (n=19 embryos, 61 cells) depletion. Orange arrowheads indicate proper actin polarization in control cells (n=11 embryos, 34 cells) while white arrowheads indicate cells where actin is strongly disorganized. **H-I** Quantification of actin orientation under *pat-4* and *unc-52* depletion, respectively. All embryos are imaged at the 2-fold stage except in B. Scale bars: 5 μm.

**Figure S2:**
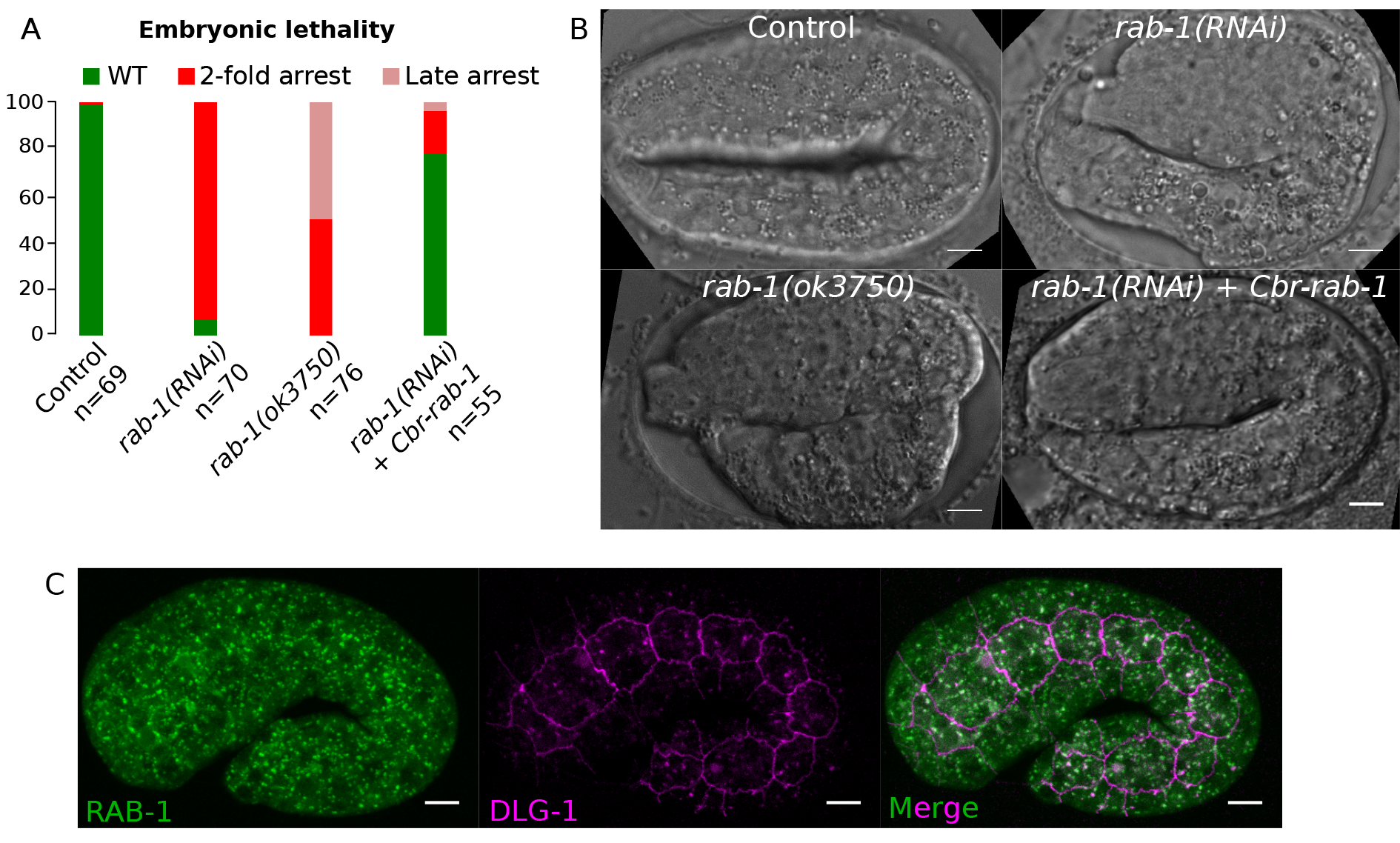
Specificity of the RNAi targeting *rab-1* and RAB-1 expression pattern. **A-B** The expression of an RNAi-resistant version of *rab-1* from *C. briggsae* almost fully rescues the lethality induced by *rab-1* depletion by RNAi and the deletion allele *rab-1(ok3750)* phenocopies *rab-1(RNAi)*. Embryos in B are imaged at the 2-fold stage. **C** RAB-1::GFP under the control of its own promoter is ubiquitously expressed in *C. elegans* embryos; the punctuate pattern observed in all epidermal cells (here in a 1.5-fold stage embryo) is consistent with an ER-Golgi function. Scale bars: 5 μm.

**Figure S3:**
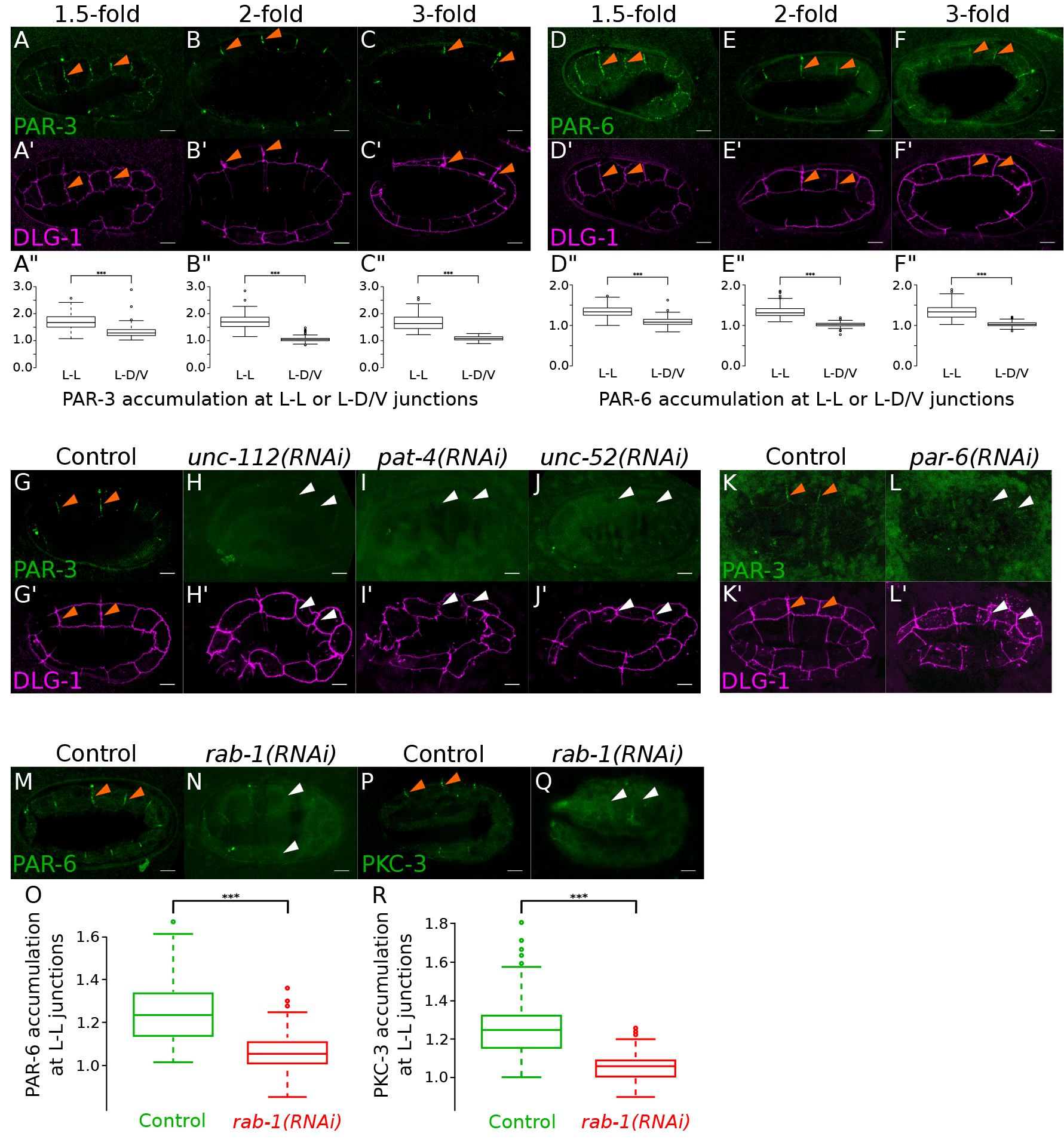
Muscles and CeHDs control PAR planar polarity in the lateral epidermis. **A-C”** Endogenous PAR-3::GFP accumulates at the lateral-lateral junctions during elongation (orange arrowheads) as revealed on the associated quantifications made on each type of junctions, lateral-lateral (L-L) and lateral-dorso-ventral (L-D/V)-embryos and numbers are the same as in Fig. 3D-G. **D-F”** Endogenous PAR-6::GFP accumulates at the L-L junctions (orange arrowheads) during elongation as revealed by the associated quantifications made on L-L and L-D/V junctions (n=10 embryos 1.5-fold, 74 L-L, 121 L-D/V; n=10 embryos 2-fold, 78 L-L, 120 L-D/V; n=10 embryos 3fold, 71 L-L, 106 L-D/V). **G-J’** Depletion of *unc-112*, *pat-4* and *unc-52* by RNAi leads to an absence of recruitment of PAR-3 at L-L junctions as depicted by white arrowheads compared to orange arrowheads in control embryos; see Fig. 3K for numbers and quantifications. **K-L’** The depletion of *par-6* by RNAi prevents PAR-3 recruitment at L-L junctions (white arrowheads), compared to control embryos where PAR-3 accumulates at L-L junctions (orange arrowheads); n=15 control embryos; n=8/11 *par-6(RNAi)* embryos. **M-R** The depletion of *rab-1* also affects the localization of endogenous PAR-6::GFP and GFP::PKC-3. In O: n=21 control embryos, 149 junctions; n=25 *rab-1(RNAi)* embryos,186 junctions. In R, n=20 control embryos, 128 junctions; n=23 *rab-1(RNAi)* embryos, 144 junctions. All embryos are imaged at the 2-fold stage except in A, C, D, F. Scale bars: 5 μm.

**Fig S4.**
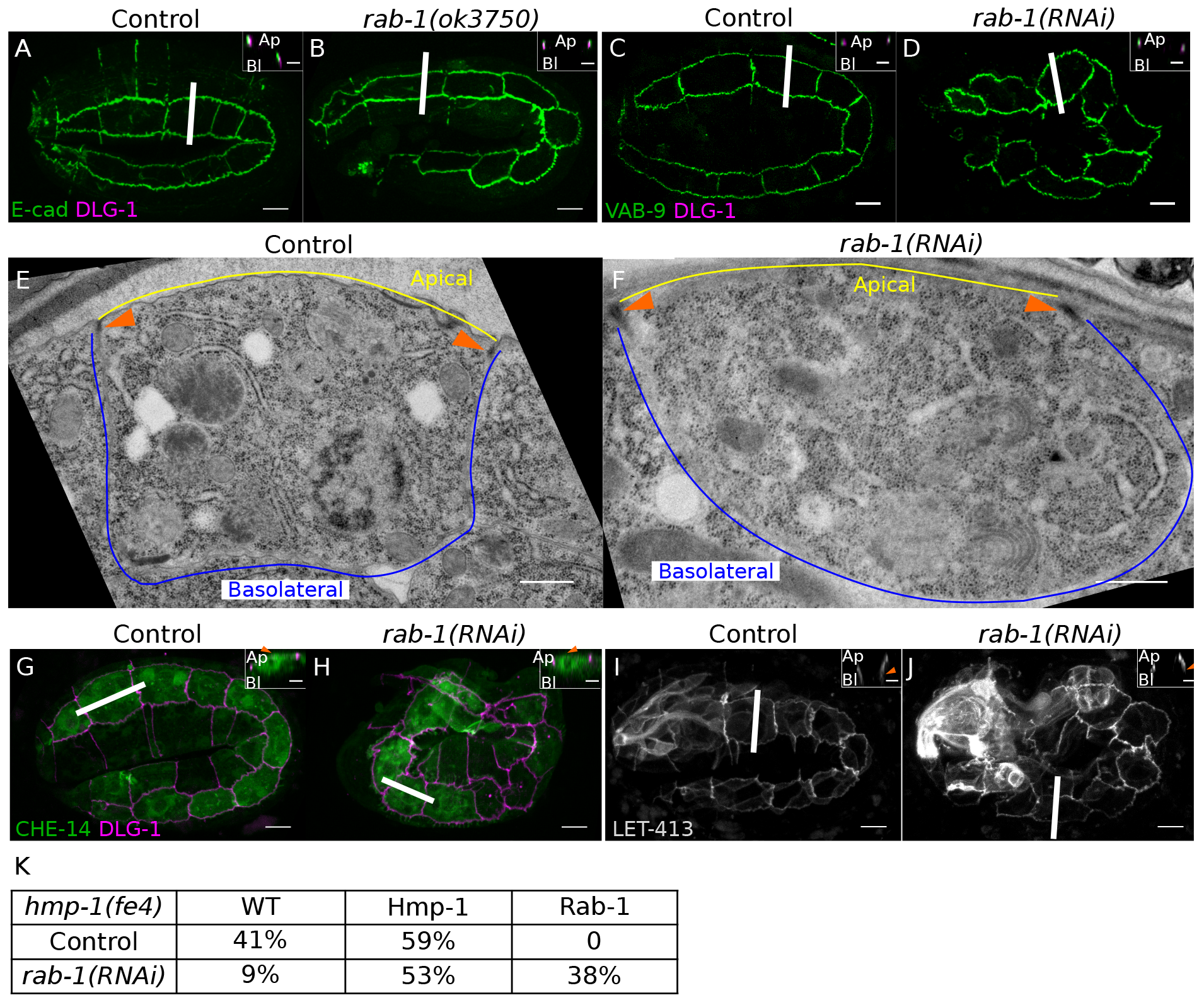
Junction integrity and apico-basal polarity are not affected upon CeHDs depletion. **A-B** As in *rab-1(RNAi)* embryos, E-cad remains above DLG-1 in *rab-1(ok3750)* mutant embryos (n=28 embryos). Small insets correspond to Z-section represented by a white line in the associated picture. **C-D** VAB-9 remains apical, above DLG-1 upon *rab-1* depletion (n=13 embryos) as in control (n=14 embryos). **E-F** Electron microscopy (EM) reveals that the electron-dense region corresponding to the junctions (orange arrowhead) is still properly localized under *rab-1* depletion. **G-J** Apico-basal polarity is not affected upon *rab-1* depletion. CHE-14 remains apical (G-H), n=16 and n=20, respectively), while LET-413 remains lateral (I-J, n=18 and n=29, respectively); orange arrowheads indicate proper localization, either apical for E-cad, CHE-14 or at the lateral membrane for LET-413. **K** Quantification showing the lack of genetic interaction between *rab-1* and the weak α-catenin mutant allele *hmp-1(fe4):* the depletion of RAB-1 in *hmp-1(fe4)* embryos does not increase the lethality induced by the *hmp-1(fe4)* allele. The Hmp-1 phenotype was scored based on the characteristic S-shape/Hmp phenotype of *hmp-1* embryos while the Rab-1 phenotype corresponds to a 2-fold arrest. All embryos are imaged at the 2-fold stage including for the EM pictures. Scale bars: 5 μm, except for small insets: 2 μm, and for E-F: 0.5 μm.

**Supplementry table 1:**
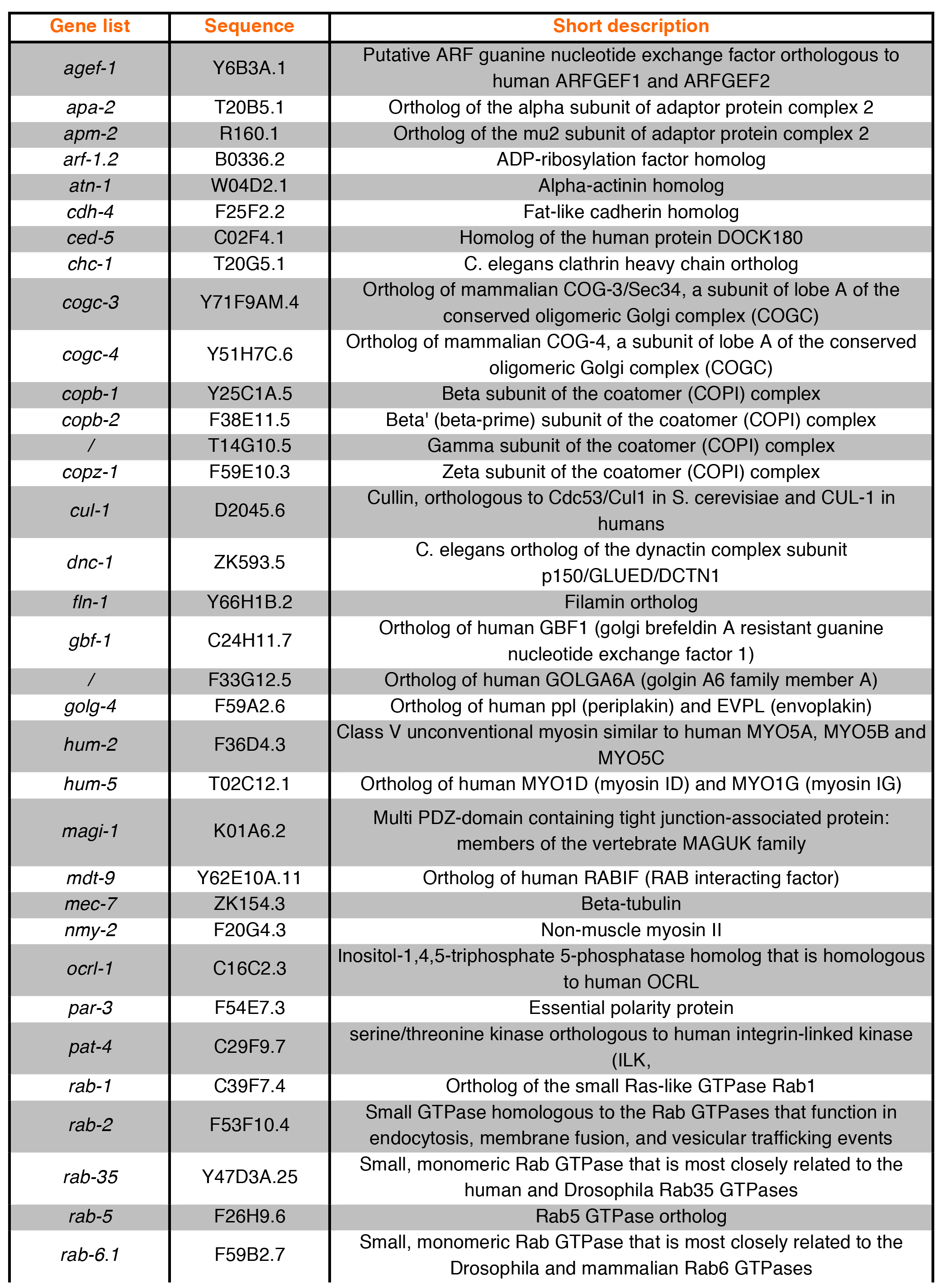
List of candidate genes targeted in the screen for actin organization.

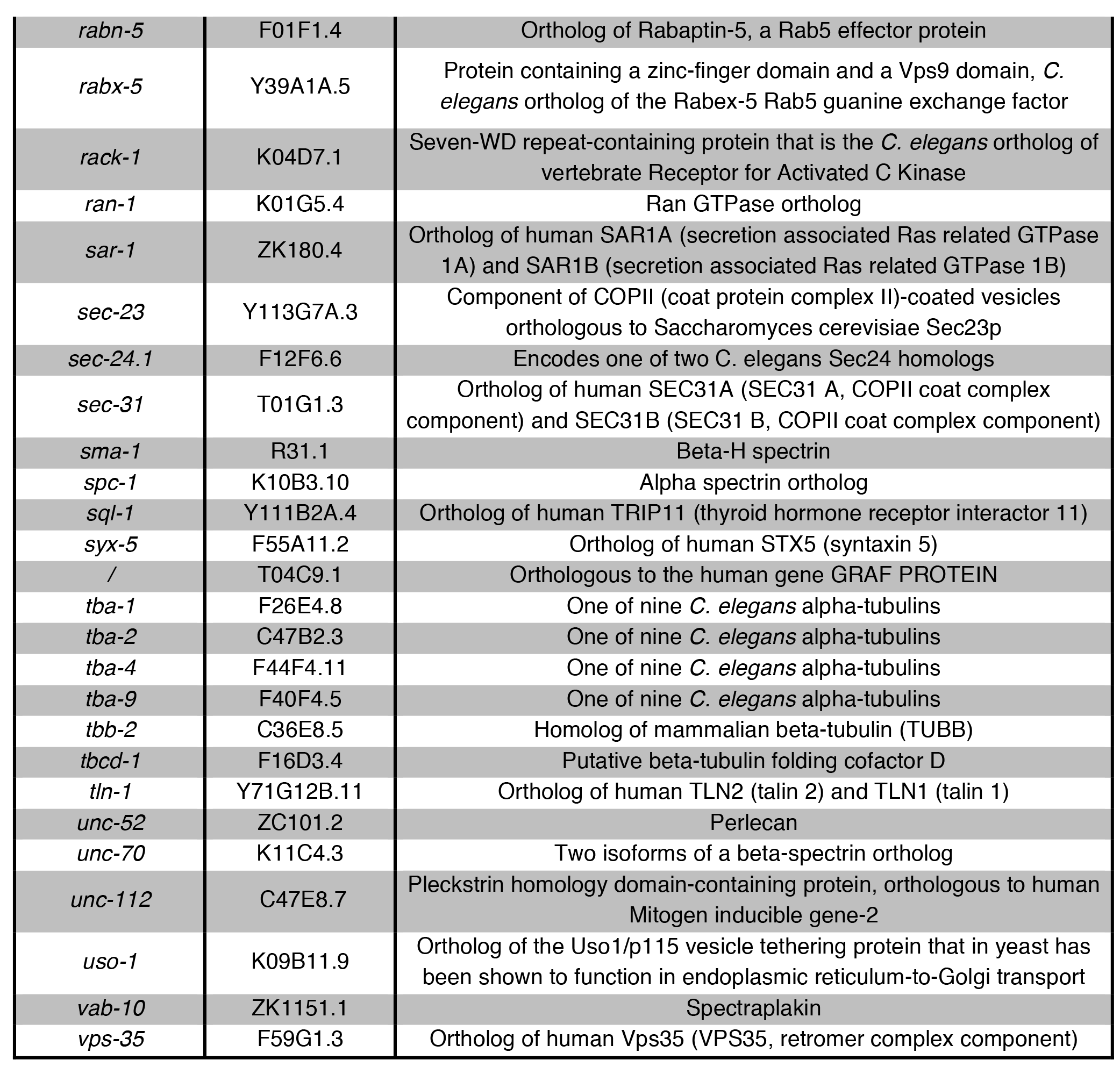

**Supplementary table 2:**
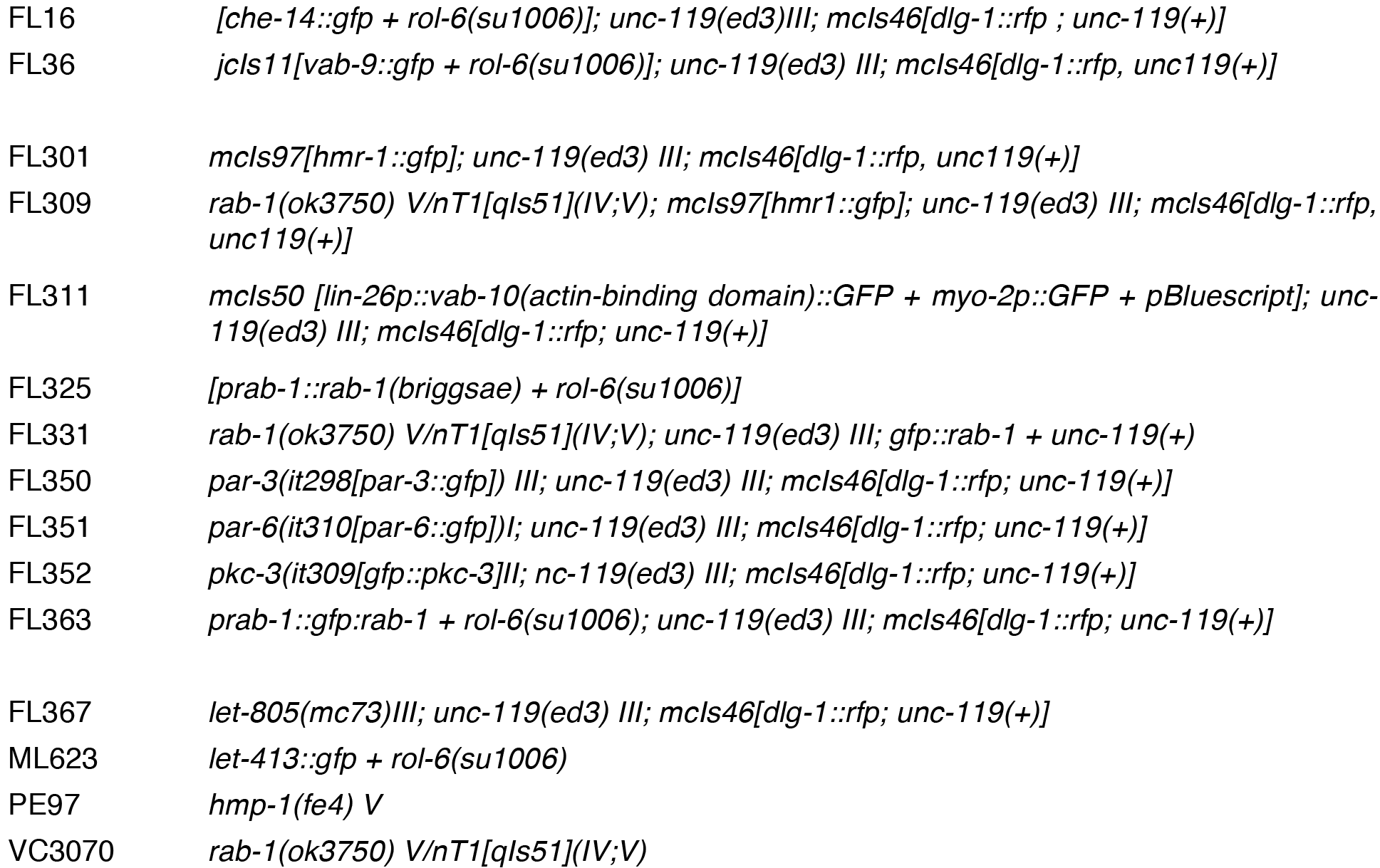
List of strains used in this study.

|  |  |
| --- | --- |
| FL16 | *[che-14::gfp + rol-6(su1006)]; unc-119(ed3)III; mcls46[dlg-1::rfp ; unc-119(+)]* |
| FL36 | *jcls11[vab-9::gfp + rol-6(su1006)]; unc-119(ed3) III; mcls46[dlg-1::rfp, unc119(+)]* |
| FL301 | *mcls97[hmr-1::gfp]; unc-119(ed3) III; mcls46[dlg-1::rfp, unc119(+)]* |
| FL309 | *rab-1(ok3750) V/nT1[qIs51](IV;V); mcls97[hmr1::gfp]; unc-119(ed3) III; mcls46[dlg-1::rfp, unc119(+)]* |
| FL311 | *mcls50 [lin-26p::vab-10(actin-binding domain)::GFP + myo-2p::GFP + pBluescript]; unc-119(ed3) III; mcls46[dlg-1::rfp; unc-119(+)]* |
| FL325 | *[prab-1::rab-1(briggsae) + rol-6(su1006)]* |
| FL331 | *rab-1(ok3750) V/nT1[qIs51](IV;V); unc-119(ed3) III; gfp::rab-1 + unc-119(+)* |
| FL350 | *par-3(it298[par-3::gfp]) III; unc-119(ed3) III; mcls46[dlg-1::rfp; unc-119(+)]* |
| FL351 | *par-6(it310[par-6::gfp])I; unc-119(ed3) III; mcls46[dlg-1::rfp; unc-119(+)]* |
| FL352 | *pkc-3(it309[gfp::pkc-3])II; nc-119(ed3) III; mcls46[dlg-1::rfp; unc-119(+)]* |
| FL363 | *prab-1::gfp::rab-1 + rol-6(su1006); unc-119(ed3) III; mcls46[dlg-1::rfp; unc-119(+)]* |
| FL367 | *let-805(mc73)III; unc-119(ed3) III; mcls46[dlg-1::rfp; unc-119(+)]* |
| ML623 | *let-413::gfp + rol-6(su1006)* |
| PE97 | *hmp-1(fe4) V* |
| VC3070 | *rab-1(ok3750) V/nT1[qIs51](IV;V)* |

## REFERENCES

Achilleos, A., Wehman, A. M. and Nance, J. (2010). PAR-3 mediates the initial clustering and apical localization of junction and polarity proteins during C. elegans intestinal epithelial cell polarization. Development 137, 1833–1842.

Aigouy, B., Farhadifar, R., Staple, D. B., Sagner, A., Roper, J. C., Julicher, F. and Eaton, S.(2010). Cell flow reorients the axis of planar polarity in the wing epithelium of Drosophila. Cell 142, 773–786.

Balklava, Z., Pant, S., Fares, H. and Grant, B. D. (2007). Genome-wide analysis identifies a general requirement for polarity proteins in endocytic traffic. Nature cell biology 9, 1066–1073.

Bosher, J. M., Hahn, B. S., Legouis, R., Sookhareea, S., Weimer, R. M., Gansmuller, A., Chisholm, A. D., Rose, A. M., Bessereau, J. L. and Labouesse, M. (2003). The Caenorhabditis elegans vab-10 spectraplakin isoforms protect the epidermis against internal and external forces. J Cell Biol 161, 757–768.

Brenner, S. (1974). The genetics of Caenorhabditis elegans. Genetics 77, 71–94.

Bulgakova, N. A., Grigoriev, I., Yap, A. S., Akhmanova, A. and Brown, N. H. (2013). Dynamic microtubules produce an asymmetric E-cadherin-Bazooka complex to maintain segment boundaries. J Cell Biol 201, 887–901.

Butler, L. C., Blanchard, G. B., Kabla, A. J., Lawrence, N. J., Welchman, D. P., Mahadevan, L., Adams, R. J. and Sanson, B. (2009). Cell shape changes indicate a role for extrinsic tensile forces in Drosophila germ-band extension. Nature cell biology 11, 859–864.

Cetera, M., Ramirez-San Juan, G. R., Oakes, P. W., Lewellyn, L., Fairchild, M. J., Tanentzapf, G., Gardel, M. L. and Horne-Badovinac, S. (2014). Epithelial rotation promotes the global alignment of contractile actin bundles during Drosophila egg chamber elongation. Nat Commun 5, 5511.

Collinet, C., Rauzi, M., Lenne, P. F. and Lecuit, T. (2015). Local and tissue-scale forces drive oriented junction growth during tissue extension. Nature cell biology 17, 1247–1258.

Costa, M., Raich, W., Agbunag, C., Leung, B., Hardin, J. and Priess, J. R. (1998). A putative catenin-cadherin system mediates morphogenesis of the Caenorhabditis elegans embryo. The Journal of cell biology 141, 297–308.

Erami, Z., Timpson, P., Yao, W., Zaidel-Bar, R. and Anderson, K. I. (2015). There are four dynamically and functionally distinct populations of E-cadherin in cell junctions. Biol Open 4, 1481–1489.

Fire, A., Xu, S., Montgomery, M. K., Kostas, S. A., Driver, S. E. and Mello, C. C. (1998). Potent and specific genetic interference by double-stranded RNA in Caenorhabditis elegans. Nature 391, 806–811.

Gillard, G., Shafaq-Zadah, M., Nicolle, O., Damaj, R., Pecreaux, J. and Michaux, G. (2015). Control of E-cadherin apical localisation and morphogenesis by a SOAP-1/AP-1/clathrin pathway in C. elegans epidermal cells. Development 142, 1684–1694.

Hresko, M. C., Schriefer, L. A., Shrimankar, P. and Waterston, R. H. (1999). Myotactin, a novel hypodermal protein involved in muscle-cell adhesion in Caenorhabditis elegans. J Cell Biol 146, 659–672.

Hubaud, A. and Pourquie, O. (2014). Signalling dynamics in vertebrate segmentation. Nature reviews. Molecular cell biology 15, 709–721.

Kamath, R. S. and Ahringer, J. (2003). Genome-wide RNAi screening in Caenorhabditis elegans. Methods 30, 313–321.

Legouis, R., Gansmuller, A., Sookhareea, S., Bosher, J. M., Baillie, D. L. and Labouesse, M. (2000). LET-413 is a basolateral protein required for the assembly of adherens junctions in Caenorhabditis elegans. Nature cell biology 2, 415–422.

Leung, B., Hermann, G. J. and Priess, J. R. (1999). Organogenesis of the Caenorhabditis elegans intestine. Developmental biology 216, 114–134.

Levayer, R. and Lecuit, T. (2013). Oscillation and polarity of E-cadherin asymmetries control actomyosin flow patterns during morphogenesis. Developmental cell 26, 162–175.

Mackinnon, A. C., Qadota, H., Norman, K. R., Moerman, D. G. and Williams, B. D. (2002). C. elegans PAT-4/ILK functions as an adaptor protein within integrin adhesion complexes. Current biology: CB 12, 787–797.

Michaux, G., Gansmuller, A., Hindelang, C. and Labouesse, M. (2000). CHE-14, a protein with a sterol-sensing domain, is required for apical sorting in C. elegans ectodermal epithelial cells. Current biology: CB 10, 1098–1107.

Olguin, P., Glavic, A. and Mlodzik, M. (2011). Intertissue mechanical stress affects Frizzled-mediated planar cell polarity in the Drosophila notum epidermis. Current biology: CB 21, 236–242.

Petit, F., Sears, K. E. and Ahituv, N. (2017). Limb development: a paradigm of gene regulation. Nat Rev Genet.

Plutner, H., Cox, A. D., Pind, S., Khosravi-Far, R., Bourne, J. R., Schwaninger, R., Der, C. J. and Balch, W. E. (1991). Rab1b regulates vesicular transport between the endoplasmic reticulum and successive Golgi compartments. The Journal of cell biology 115, 31–43.

Rauzi, M., Lenne, P. F. and Lecuit, T. (2010). Planar polarized actomyosin contractile flows control epithelial junction remodelling. Nature 468, 1110–1114.

Rogalski, T. M., Mullen, G. P., Gilbert, M. M., Williams, B. D. and Moerman, D. G. (2000). The UNC-112 gene in Caenorhabditis elegans encodes a novel component of cell-matrix adhesion structures required for integrin localization in the muscle cell membrane. J Cell Biol 150, 253–264.

Rogalski, T. M., Williams, B. D., Mullen, G. P. and Moerman, D. G. (1993). Products of the unc-52 gene in Caenorhabditis elegans are homologous to the core protein of the mammalian basement membrane heparan sulfate proteoglycan. Genes & development 7, 1471–1484.

Sagner, A., Merkel, M., Aigouy, B., Gaebel, J., Brankatschk, M., Julicher, F. and Eaton, S. (2012). Establishment of global patterns of planar polarity during growth of the Drosophila wing epithelium. Current biology: CB 22, 1296–1301.

Schmid, T. and Hajnal, A. (2015). Signal transduction during C. elegans vulval development: a NeverEnding story. Current opinion in genetics & development 32, 1–9.

Segev, N., Mulholland, J. and Botstein, D. (1988). The yeast GTP-binding YPT1 protein and a mammalian counterpart are associated with the secretion machinery. Cell 52, 915–924.

Shafaq-Zadah, M., Brocard, L., Solari, F. and Michaux, G. (2012). AP-1 is required for the maintenance of apico-basal polarity in the C. elegans intestine. Development 139, 2061–2070.

Shyer, A. E., Rodrigues, A. R., Schroeder, G. G., Kassianidou, E., Kumar, S. and Harland, R. M. (2017). Emergent cellular self-organization and mechanosensation initiate follicle pattern in the avian skin. Science 357, 811–815.

Simoes Sde, M., Blankenship, J. T., Weitz, O., Farrell, D. L., Tamada, M., Fernandez-Gonzalez, R. and Zallen, J. A. (2010). Rho-kinase directs Bazooka/Par-3 planar polarity during Drosophila axis elongation. Developmental cell 19, 377–388.

Simske, J. S., Koppen, M., Sims, P., Hodgkin, J., Yonkof, A. and Hardin, J. (2003). The cell junction protein VAB-9 regulates adhesion and epidermal morphology in C. elegans. Nature cell biology 5, 619–625.

Vuong-Brender, T. T., Ben Amar, M., Pontabry, J. and Labouesse, M. (2017). The interplay of stiffness and force anisotropies drives embryo elongation. Elife 6.

Vuong-Brender, T. T., Yang, X. and Labouesse, M. (2016). C. elegans Embryonic Morphogenesis. Curr Top Dev Biol 116, 597–616.

Zahreddine, H., Zhang, H., Diogon, M., Nagamatsu, Y. and Labouesse, M. (2010). CRT-1/calreticulin and the E3 ligase EEL-1/HUWE1 control hemidesmosome maturation in C. elegans development. Current biology: CB 20, 322–327.

Zhang, H., Landmann, F., Zahreddine, H., Rodriguez, D., Koch, M. and Labouesse, M. (2011). A tension-induced mechanotransduction pathway promotes epithelial morphogenesis. Nature 471, 99–103.

